# Vaporized cannabis extracts have reinforcing properties and support conditioned drug-seeking behavior in rats

**DOI:** 10.1101/791319

**Authors:** Timothy G. Freels, Lydia N. Baxter-Potter, Janelle M. Lugo, Nicholas C. Glodosky, Hayden R. Wright, Samantha L. Baglot, Gavin N. Petrie, Z Yu, Brian H. Clowers, Carrie Cuttler, Rita A. Fuchs, Matthew N. Hill, Ryan J. McLaughlin

## Abstract

Recent trends in cannabis legalization have increased the necessity to better understand the effects of cannabis use. Animal models involving traditional cannabinoid self-administration approaches have been notoriously difficult to establish and differences in the drug employed and its route of administration have limited the translational value of preclinical studies. To address this challenge in the field, we have developed a novel method of cannabis self-administration using response-contingent delivery of vaporized Δ_9_-tetrahydrocannabinol-rich (CAN_THC_) or cannabidiol-rich (CAN_CBD_) complete cannabis extracts. Male Sprague Dawley rats were trained to nosepoke for discrete puffs of CAN_THC_, CAN_CBD_, or vehicle (VEH) in daily one-hour sessions. Cannabis vapor reinforcement resulted in strong discrimination between active and inactive operanda. CAN_THC_ maintained higher response rates under fixed ratio schedules and higher break points under progressive ratio schedules compared to CAN_CBD_ or VEH, and the number of vapor deliveries positively correlated with plasma THC concentrations. Moreover, metabolic phenotyping studies revealed alterations in locomotor activity, energy expenditure, and daily food intake that are consistent with effects in human cannabis users. Furthermore, both cannabis regimens produced ecologically relevant brain concentrations of THC and CBD and CAN_THC_ administration decreased hippocampal CB1 receptor binding. Removal of CAN_THC_ reinforcement (but not CAN_CBD_) resulted in a robust extinction burst and an increase in cue-induced cannabis-seeking behavior relative to VEH. These data indicate that volitional exposure to THC-rich cannabis vapor has *bona fide* reinforcing properties and collectively support the utility of the vapor self-administration model for the preclinical assessment of volitional cannabis intake and cannabis-seeking behaviors.

## INTRODUCTION

With several states recently passing legislation allowing for the use of cannabis for recreational purposes, there is a pressing need to better understand the neurobiological effects of cannabis use/misuse. Animal models are valuable in this regard because they afford precise control over extraneous variables that complicate the interpretation of cross-sectional human data. However, current approaches have limited translational value [1]. Synthetic cannabinoid receptor 1 (CB1R) agonists or isolated cannabis constituents (e.g., Δ_9_-tetrahydrocannabinol [THC], cannabidiol [CBD]) have become the drug of choice in rodent models of cannabis exposure; even though the pharmacological properties of these compounds differ greatly from those of inhaled cannabis. THC and synthetic CB1R agonists have different pharmacological profiles (partial vs. full agonists) and recruit different intracellular signaling pathways [2]. Thus, synthetic CB1R agonists may fail to recapitulate the effects of THC, let alone the effects of cannabis.

Animal models that involve access to THC alone have drawbacks as well. Over 120 unique phytocannabinoid compounds are present in *Cannabis sativa* in addition to THC. These phytocannabinoids have distinct pharmacological and behavioral profiles [3], and interactions between phytocannabinoids give rise to the psychotropic or physiological effects of cannabis chemovars [4–6]. For example, CBD acts as a CB1R negative allosteric modulator and inhibits THC-dependent intracellular signaling and beta-arrestin-2-mediated CB1R internalization [7,8]. CBD attenuates THC-induced paranoia and memory impairments in humans [9] but can also increase THC serum concentrations when co-administered with THC [10]. Therefore, THC administration alone may not produce effects that are representative of the effects of cannabis exposure in humans. In line with this, pure THC self-administration has been difficult to demonstrate at the preclinical level [11]. Rats acquire stable rates of intravenous THC:CBD (10:1) self-administration only after extended passive pre-exposure to vaporized THC:CBD [12]. Thus, studying the effects of THC in the presence of other phytocannabinoids is important for modeling cannabis use.

Current animal models typically use the intravenous route for drug delivery; even though, the most common route of recreational cannabis use is inhalation [13], and the pharmacokinetics of cannabinoids vary considerably depending on administration route [14–16]. For example, intravenous THC administration produces adverse effects in humans, likely related to high dosing and faster infusion rates [17]. Similarly, vaporized THC administration produces conditioned place preference, whereas intraperitoneal THC administration produces conditioned place avoidance in rodents [18]. Thus, the route of cannabis administration may fundamentally influence the degree to which cannabis can support self-administration.

A more translationally relevant approach is needed that uses cannabis and mimics the most common route of administration in human users. With this in mind, we have developed a novel, ecologically valid model of cannabis vapor self-administration that uses ‘e-cigarette’ technology to deliver discrete puffs of vaporized cannabis extracts to rodents in a response-contingent manner. We used this procedure to examine whether vaporized cannabis extracts that are high in THC or CBD have reinforcing properties that support stable drug-taking behavior. We characterized the metabolic phenotype of cannabis-trained rats and since human cannabis users and rodents chronically treated with THC exhibit reduced CB1R availability [19] and decreased CB1R expression [20–23], respectively, we also measured hippocampal CB1 receptor binding in rats from each treatment group. Furthermore, given that acute abstinence can unmask withdrawal-related affective symptoms for other drugs of abuse [24], we examined whether acute forced abstinence from cannabis vapor increases anxiety-like behavior. Finally, we tested whether vaporized cannabis regimens maintain drug-seeking behavior in the drug-predictive context (i.e., under extinction conditions) or upon response-contingent presentation of a cannabis-paired light stimulus (i.e., cue-induced reinstatement).

## METHODS AND MATERIALS

A full description of all experimental procedures is provided in the **Supplemental Materials**.

### Animals

Male Sprague-Dawley rats (Simonsen Laboratories, Gilroy, CA; 350-400 g) were pair-housed in a humidity-controlled vivarium on a 12:12 reverse light-dark schedule (lights off at 7h00). Food and water were available *ad libitum*. All procedures followed the National Institutes of Health Guide for the Care and Use of Laboratory Animals and were approved by the Washington State University Institutional Animal Care and Use Committee.

### Drugs

Cannabis extracts rich in THC (CAN_THC_; 28.4% THC/1.1% CBD) or CBD (CAN_CBD_; 1.96% THC/59.3% CBD) were heated to 60°C under constant stirring and dissolved in 80% propylene glycol/20% vegetable glycerol vehicle (VEH) at a concentration of 400 mg/ml based on previous studies [25,26]. Based on the certificate of analysis provided by NIDA, the final concentrations of THC and CBD in CAN_THC_ were ~116.8 mg/ml and ~4.4 mg/ml, respectively. The final concentrations of THC and CBD in CAN_CBD_ were ~7.84 mg/ml and 237.2 mg/ml. The CB1R antagonist, AM251 (Cayman Chemical; Ann Arbor, MI), was dissolved in dimethyl sulfoxide:Tween-80:saline (1:1:18) vehicle and administered at a dose of 0, 1, or 3 mg/kg (1 ml/kg, i.p.).

### Self-Administration Training

Rats were trained to nosepoke for CAN_THC_, CAN_CBD_, or VEH vapor puffs under a fixed ratio-1 (FR-1) reinforcement schedule during daily one-hour sessions on 11 consecutive days (see **Supplement** for a description of the vapor delivery system). Rats then progressed to an FR-2 schedule (days 12-16) and then to an FR-4 schedule (day 17-21). Nosepoke responses made on one (active) operandum resulted in a 3-s activation of the vaporizer and illumination of a cue light. The cue light remained illuminated during a 60-s time-out period, during which responses were not reinforced. Nosepoke responses made on the other (inactive) operandum had no programmed consequences. On the final self-administration day (day 22), responding was reinforced under a progressive ratio (PR) schedule for 180-min. Schedule demand increased after each vapor delivery according to the following schedule: 1, 1, 2, 2, 3, 3, 4, 4, 5, 5, 7, 7, 9, 9, 11, 11, 13, 13, 15, 15, 18, 18, 21, 21, 24, 24, etc. [27]. The breakpoint was defined as the total number of vapor deliveries obtained until responding ceased for a minimum of 15 minutes.

### Cannabinoid Quantification

Immediately following a single one-hour CAN_THC_ or CAN_CBD_ (200 or 400 mg/ml) self-administration session, blood samples (~100 μl) were collected in a different cohort of rats (N=24) to evaluate the relationship between the total number of vapor deliveries and circulating THC and CBD concentrations. Blood was collected vial the tail vein in sterile tubes containing 10 ml ethylenediaminetetraacetic acid, centrifuged at 4°C at 4,000 g for 15 min, and stored at −20°C. Another subset of rats (N=21) was sacrificed 24 hr after the final self-administration session. Brains were extracted to analyze THC and CBD tissue levels. THC and CBD concentrations in plasma and brain were quantified as described previously [10,28] (see **Supplement**).

### CB1R Radioligand Binding Assay

Frozen whole brain tissue was collected from some rats (N=12) 24 hr after the final self-administration session. Tissue was homogenized in TME buffer and centrifuged to generate the crude membrane fraction. Protein concentrations were determined using the Bradford method (Bio-Rad). Membranes (10-μg protein/sample) were incubated in TME buffer with [_3_H]CP55,940 (0.25, 0.5, 1.25, or 2.5 nM) in the absence or presence of AM251 (10 μM) to assess total and nonspecific binding, respectively. *B*_max_ (maximal binding site density) and *K*_d_ (binding affinity) were calculated by nonlinear curve fitting to the single site Michaelis–Menten equation using GraphPad Prism as described previously [29,30].

### Radiotelemetry Recordings

A subset of rats (N=8) was anesthetized with isoflurane/oxygen mixture (isoflurane 5% induction, 1-3% maintenance). Sterile radiotelemetry transmitters (Starr Life Sciences Corp., Oakmont, PA; PTD 4000) were implanted (see **Supplement**). Radiotelemetry transmissions indexing locomotor activity and body temperature were collected daily during the one-hour self-administration sessions by Respironics ER-4000 receiver plates placed under the chambers.

### Metabolic Phenotyping

A subset of rats (N=18) were housed individually in metabolic cages, when not in the self-administration chambers. Feeding behavior, water intake, energy expenditure, physical activity, and respiratory quotient (RQ) were continuously monitored (see **Supplement**).

### CB1R Antagonism

Rats (N=16) were trained to self-administer vaporized CAN_THC_ or CAN_CBD_ (400 mg/ml) under an FR-1 schedule over 26 daily sessions. Rats received mock IP injections on the two days prior to testing the effects of systemic CB1R antagonism on training days 16, 21, and 26. One hour before testing, rats received AM251 (1 or 3 mg/kg, IP) or VEH using a counterbalanced within-subjects design. Additional training sessions were conducted between test sessions to re-establish stable responding. All data were converted to a percent change score relative to the previous mock injection day.

### Elevated Plus Maze Test

Anxiety-like behavior was assessed in another cohort of rats (N=37) 24 hours after the final self-administration session using the elevated plus maze (EPM) test as described previously [30,31] (see **Supplement**).

### Extinction Training and Cue-Induced Reinstatement

Rats (N=35) were trained to self-administer CAN_THC_, CAN_CBD_, or VEH under an FR-1 schedule over 19 daily sessions. Rats then underwent daily one-hour extinction training sessions during which nosepoke responses had no programmed consequences. Extinction training continued until rats reached the extinction criterion (i.e., ≥ seven extinction training sessions with ≥50% decrease in active nosepoke responses during the final two sessions). Rats were then tested for cue-induced reinstatement of extinguished drug-seeking behavior. During the one-hour test session, active nosepoke responses resulted in cue light presentations without vapor delivery.

### Statistical Analysis

Vapor deliveries, active and inactive responses, trials to extinction, nosepoke discrimination, body weight, within-session locomotor activity, food intake, water intake, lounge time, locomotor activity, energy expenditure, respiratory quotient, and EPM data were compared across groups using planned t-tests or one-way or mixed factor analyses of variance (ANOVA) with treatment group (CAN_THC_, CAN_CBD_, VEH) as the between-subjects factor and time (session, 15-min bin, phase) as the within-subjects factor. Nosepoke discrimination was calculated based on a formula used in [12]: nosepoke discrimination index = (active nosepokes – inactive nosepokes)/(active nosepokes + inactive nosepokes), where 0 indicated no discrimination between active and inactive operanda, 1 indicated complete preference for the active operandum, and −1 indicated complete preference for inactive operandum [12]. Effects of AM251 were analyzed using separate repeated-measures one-way ANOVAs. Significant effects were further probed using Tukey’s or Dunnett’s post-hoc tests. Vapor delivery and plasma cannabinoid concentrations were correlated using Pearson’s correlation coefficient (*r*). Alpha was set at .05. Effect sizes are reported as 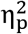.

## RESULTS

### THC-Rich Cannabis Vapor Supports Stable Rates of Self-Administration

CAN_THC_ elicited more active responses compared to VEH on days 13, 14, 16-2 1 and compared to CAN_CBD_ on days 5, 6, 9-21 (interaction: *F*_(40,540)_=3.57, *p*=.001, 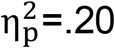, post-hoc *p*’s<.05, **Fig. 1C**), predominantly under the more demanding FR-2 and FR-4 reinforcement schedules. CAN_CBD_ elicited fewer active responses compared to VEH on days 1 and 2 (*p*’s=.01 and .03, respectively). CAN_CBD_ also elicited fewer inactive responses than CAN_THC_ and VEH (main effects:*F*_(2-20, 27-540)_=6.27–8.36, *p’s* ≤.001, 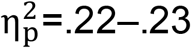, post-hoc *p*’s≤.03).

**Figure 1.**
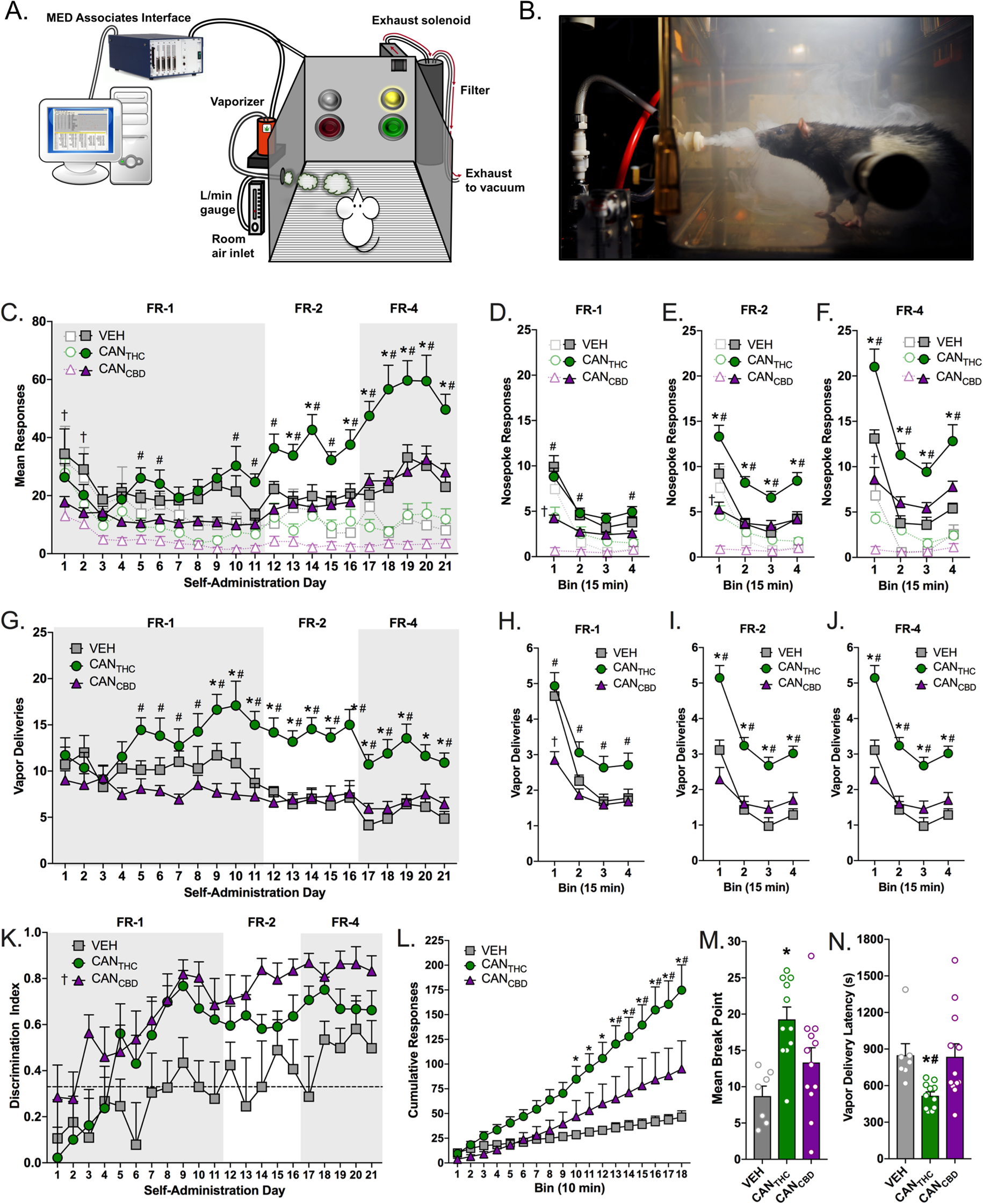
Cannabis vapor has reinforcing properties and supports stable rates of self-administration in male rats. **(A)** Schematic illustration of the vapor self-administration apparatus, (adapted from [47)], and **(B)** real-life depiction of a rat responding for cannabis vapor. **(C)** Mean active (colored symbols) and inactive (open symbols) nosepoke responding for vapor containing high concentrations of THC (CAN_THC_), CBD (CAN_CBD_), or vehicle (VEH) across increasing fixed ratio schedules of reinforcement. **(D-F)** Mean active (colored symbols) and inactive (open symbols) nosepoke responding organized by 15 min bins within **(D)** FR-1, **(E)** FR-2, and **(F)** FR-4 schedules of reinforcement. **(G)** Mean number of CAN_THC_, CAN_CBD_, or VEH vapor deliveries earned across increasing fixed ratio schedules of reinforcement. **(H-J)** Mean number of vapor deliveries earned organized by 15 min binds within **(H)** FR-1, **(I)** FR-2, and **(J)** FR-4 schedules of reinforcement. **(K)** Nosepoke operanda discrimination index for CAN_THC_, CAN_CBD_, and VEH vapor across increasing fixed schedules of reinforcement. The dotted line represents a discrimination index of 0.33, which indicates a 2:1 rate of active:inactive nosepoke responding. **(L)** Mean cumulative number of active responses for CAN_THC_, CAN_CBD_, and VEH vapor during a 3 hr progressive ratio challenge. Data are tallied and organized into 10 min bins. **(M)** Mean break points for CAN_THC_, CAN_CBD_, and VEH vapor during the progressive ration challenge (defined as an absence of active nosepoke responding for period of 15 min. **(N)** Mean latency to initiate active nosepoke responding for CAN_THC_, CAN_CBD_, or VEH vapor relative to the immediately preceding vapor delivery. n=7-12/group, *p* ≤ .05. * denotes significant differences between CAN_THC_ and VEH groups. # denotes significant differences between CAN_THC_ and CAN_CBD_ groups. † denotes significant differences between CAN_CBD_ and VEH groups.

The majority of responding occurred during the first 15 min of the session under each reinforcement schedule (**Figs. 1D-F**). However, CAN_THC_ elicited more active responses than VEH and CAN_CBD_ under the FR-2 (**Fig. 1E**) and FR-4 (**Fig. 1F**) schedules during each 15-min bin (interactions: *F*_(6,81)_=6.68–9.05, *p*’s<.001, 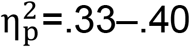, post-hoc *p*’s ≤.005). Conversely, CAN_CBD_ elicited fewer active responses than VEH during the first 15-min bin under each reinforcement schedule (*p*’s ≤.037). CAN_CBD_ elicited less inactive responding than CAN_THC,_ which in turn elicited less inactive lever responding than VEH during the first 15-min of sessions under each reinforcement schedule (interactions: *F*_(6,81)_=6.26–9.47, *p*’s<.001, 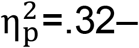 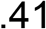, post-hoc *p*’s ≤.02) (**Figs. 1D-F**), but CAN_THC_ elicited more inactive responses than VEH (*p*=.04) and CAN_CBD_ (*p*=.01) during the second 15-min of sessions under the FR-4 schedule (**Fig. 1F**).

Stable rates of vapor delivery were achieved under each reinforcement schedule, as indicated by a lack of day effect over the final three days under each schedule. CAN_THC_ elicited more vapor deliveries than VEH on days 9-21 and more than CAN_CBD_ on days 5-19 and 21 (*F*_(40,540)_=2.24, *p*=.001, 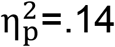, post-hoc *p*’s<.05, **Fig. 1G**). CAN_THC_ elicited more vapor deliveries than VEH under FR-2 and FR-4 schedules and more than CAN_CBD_ under each reinforcement schedule (interactions: *F*_(6,81)_=7.91–12.3, *p*’s<.001, 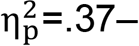 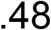, post-hoc *p*’s≤.01, **Figs. 1H-J**). CAN_CBD_ elicited fewer vapor deliveries than VEH during for first 15-min bin under the FR-1 schedule (*p*<.001) (**Fig. 1H**).

The average nosepoke discrimination index was significantly above 0.33 for CAN_THC_ (*t*_(20)_=4.39, *p*<.001) and CAN_CBD_ (*t*_(20)_=8.77, *p*<.001), but not for VEH. Thus, only the cannabis vapor self-administering groups obtained active to inactive responding ratios that were greater than 2:1. Furthermore, CAN_CBD_ elicited better discrimination for the active operandum than VEH (*F*_(2,27)_=6.57, *p*=.001, 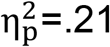 post-hoc *p*=.003, **Fig. 1K**).

### THC-Rich Cannabis Vapor Exhibits Robust Motivational Properties

During the PR schedule of reinforcement, CAN_THC_ produced higher cumulative responding compared to VEH from 100-120 min and compared to both VEH and CAN_CBD_ from 130-180 min (*F*_(36,486)_=4.49, *p*=.001, 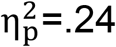, post-hoc *p*’s≤.04, **Fig. 1L**). CAN_THC_ also elicited higher break points than VEH (*F*_(2,27)_=7.17, *p*=.001, 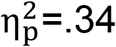, post-hoc *p*=.001) and a trend for higher break points than CAN_CBD_ (*p*=.058, **Fig. 1M**). Moreover, CAN_THC_ elicited shorter latencies to re-initiate responding following vapor deliveries than VEH and CAN_CBD_ (*F*_(2,27)_=5.01, *p*=.01, 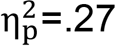, post-hoc *p*’s≤.04, **Fig. 1N**). Importantly, PR responding for CAN_CBD_ was not different from VEH on any endpoint.

### Cannabis Vapor Self-Administration Produces Biologically Relevant Increases in Plasma THC and CBD Concentrations

The number of cannabis vapor deliveries (200 or 400 mg/ml) during the one-hour self-administration session positively correlated with plasma **(A)** THC (CAN_THC-200_: *r* =.51, *p*=.03; CAN_THC-400_: *r* =.86, *p*<.001) and **(B)** CBD (CAN_CBD-200_: *r* =.58, *p*=.01; CAN_CBD-400_: *r* =.51, *p*=.18) concentrations post session. Brain THC concentrations did not differ 24 hours after the final CAN_THC_ versus CAN_CBD_ self-administration session (**Fig. 2C**), while brain tissue concentrations of CBD were higher following the CAN_CBD_ regimen compared to the CAN_THC_ regimen (*t*_(20)_=2.24, *p*=.04, 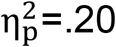, **Fig. 2D**).

**Figure 2.**
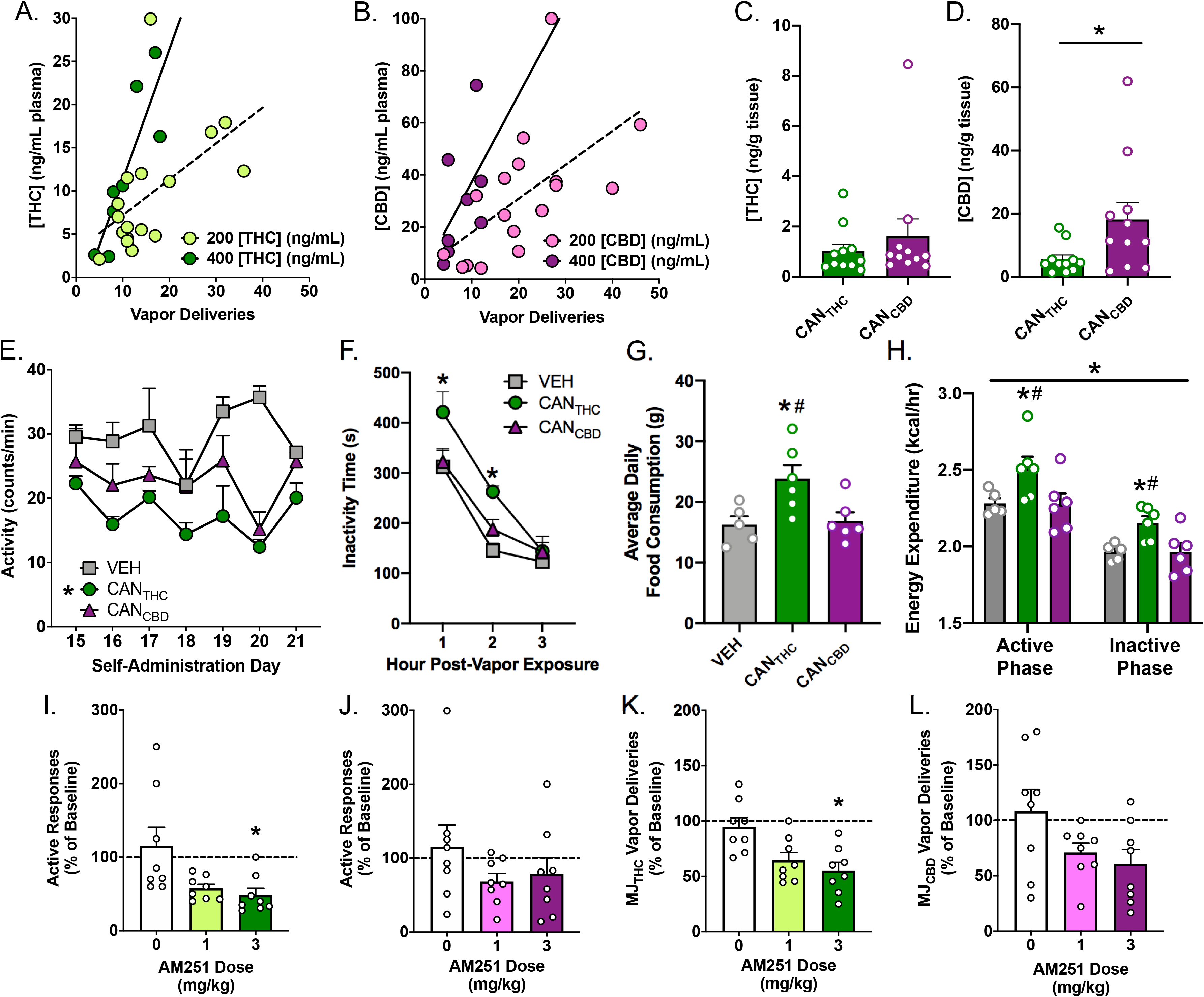
Cannabis vapor self-administration produces physiologically and behaviorally relevant plasma cannabinoid concentrations. Correlations between the number of cannabis vapor deliveries (200 or 400 mg/ml) earned and plasma concentrations of **(A)** THC (CAN_THC-200_: *r* = .51, *p* = .03; CAN_THC-400_: *r* = .86, *p* < .001) and **(B)** CBD (CAN_CBD-200_: *r* = .58, *p* = .01; CAN_CBD-400_: *r* = .51, *p* = .18) at the end of the 1-hr self-administration session. Brain tissue concentration of **(C)** THC and **(D)** CBD measured 24 hours after the final self-administration session in rats trained to self-administer CAN_THC_ or CAN_CBD_ vapor. **(E)** Radiotelemetry recordings of within-session locomotor activity (counts/min) over the final 7 days of self-administration in a subset of CAN_THC_, CAN_CBD_, and VEH self-administering rats. **(F)** Home cage inactivity (i.e., short lounging behavior) measured in the 3 hr immediately following CAN_THC_, CAN_CBD_, or VEH vapor self-administration (averaged over the final 10 days of training). **(G)** Mean total daily food consumption (g) in a subset of rats trained to self-administer CAN_THC_, CAN_CBD_, or VEH vapor (averaged over the final 10 days of training). **(H)** Mean energy expenditure (kcal/hr) during active and inactive phases of rats trained to self-administer CAN_THC_, CAN_CBD_, or VEH vapor (averaged over the final 10 days of training). **(I-J)** Mean active nosepoke responses for **(I)** CAN_THC_ and **(J)** CAN_CBD_ vapor following systemic administration of the CB1R antagonist AM251 (0, 1, or 3 mg/kg, ip). Data are depicted as a percentage of baseline from the preceding mock injection day. **(K-L)** Mean vapor deliveries earned for **(I)** CAN_THC_ and **(J)** CAN_CBD_ vapor following systemic administration of the CB1R antagonist AM251 (0, 1, or 3 mg/kg, ip). Data are depicted as a percentage of baseline from the preceding mock injection day. *p* ≤ .05. * denotes significant differences between CAN_THC_ and VEH groups. # denotes significant differences between CAN_THC_ and CAN_CBD_ groups.

### CAN_THC_ Vapor Self-Administration Alters Physical Activity and Daily Food Intake

Self-administration of CAN_THC_ (but not CAN_CBD_) reduced locomotor activity during the final seven self-administration sessions relative to VEH (*p*=0.009, **Fig. 2E**). The number of vapor deliveries negatively correlated with locomotor activity on days 13, 16, and 18 (*r’s* =.73-98; *p*’s<.05)]. CAN_THC_ reduced locomotion during the self-administration sessions relative to VEH (treatment: *F*_(2,14)_=10.2, *p*=.002; time: *F*_(2,28)_=51.2, *p*<.001; interaction: *F*_(4,28)_=1.37, *p*=.27), relative to CAN_CBD_ during the first hour post-vapor exposure, and relative to VEH during the first and second hour post-vapor exposure (*p*’s<.02, **Fig. 2F**). CAN_THC_ increased daily food consumption over the last 10 days of training relative to VEH and CAN_CBD_ (*F*_(2,14)_=5.73, *p*=.02, 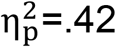, post-hoc *p*’s<.03, **Fig. 2G**), but neither CAN_THC_ nor CAN_CBD_ altered body weight gain. CAN_THC_ also increased energy expenditure compared to VEH and CAN_CBD_ (phase: *F*_(1,14)_=179.00, *p*<.001, 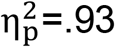; treatment: *F*_(2,14)_=5.17, *p*=.021, 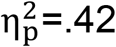, post-hoc *p*’s<.05, **Fig. 2H**) and produced higher VO_2_ and VCO_2_ values compared to CAN_CBD_ (phase: *F*_(1,14)_=98.8-140.00, *p*’s<.001, 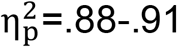; treatment: *F*_(1,14)_=4.04-4.49, *p*’s=.03, 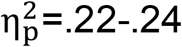. *p*’s<.05, **Fig. S1D-E**). Food intake, water intake, and distance travelled were higher during the active vs. inactive phase (*F*_(1,14)_=25.00-166, *p*’s<.001, 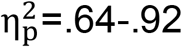) independent of treatment, and there were no effects of phase or treatment on respiratory quotients (RQ) (**Fig. S1**).

### The Reinforcing Effects of Vaporized CANTHC Require CB1 Receptor Stimulation

Systemic CB1R antagonism differentially impaired the reinforcing effects of vaporized CAN_THC_ and CAN_CBD_. Baseline active responses and vapor deliveries were not significantly different between conditions, and VEH treatment did not alter these measures relative to mock injection. The 3 mg/kg dose (but not 1 mg/kg dose) of AM251 decreased CAN_THC_-reinforced active responses (*p*=.01) and vapor deliveries (*p*=.01) relative to baseline (*F*_(2,7)_=6.05-7.44, *p*’s≤.04, 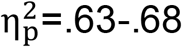, **Figs. 2I-J**). In contrast, AM251 did not significantly alter CAN_CBD_-reinforced active responses or vapor deliveries earned (**Figs. 2K-L**). AM251 treatment did not alter inactive responding relative to baseline (**Fig. S2**).

### CAN_THC_ Self-Administration Alters Hippocampal CB1 Receptor Binding

CB1R radioligand binding assays on hippocampal tissue from cannabis-exposed rats revealed that the CAN_THC_ regimen significantly reduced CB1R maximal binding site density (B_max_; *t*_(6)_=2.90, *p*=.03, ᶯ_2_=.58, **Fig. 3A**) without altering CB1R binding affinity (K_D_; *t*_(6)_=.53, *p*=.61, η_2_=.05, **Fig. 3B**) compared to VEH, whereas CAN_CBD_ failed to alter these endpoints when measured 24 hr after the final vapor self-administration session.

**Figure 3.**
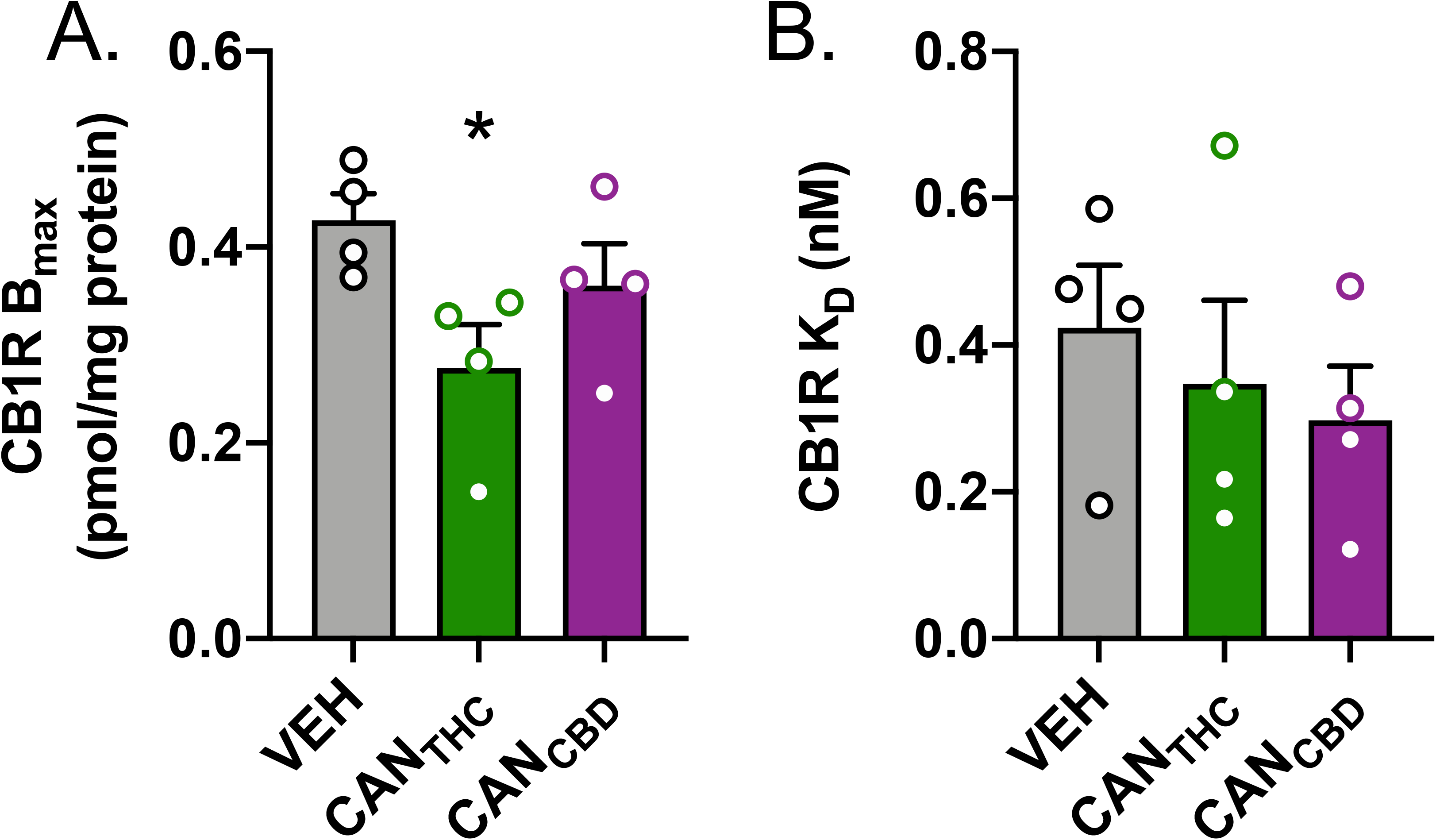
THC-rich cannabis vapor self-administration alters the maximal binding site density of CB1 receptors in the dorsal hippocampus. **(A)** hippocampal CB1R binding site density (pmol/mg protein) and **(B)** CB1R binding affinity (nM) in rats trained to self-administer CAN_THC_, CAN_CBD_, or VEH. n=4/group, *p* ≤ .05, * denotes significant differences between CAN_THC_ and VEH groups.

### Removal of Cannabis Vapor Reinforcement Elicits an Extinction Burst

Rates of vapor self-administration did not differ between groups on the final self-administration day (**Fig. S3**). Removal of CAN_THC_ or CAN_CBD_ increased active responses on the first extinction day relative to the last self-administration day (*F(*_2,32)_=3.47, *p*=0.04, 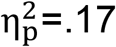; *p’*s<.005), whereas removal of VEH did not alter active responses (**Fig. 4A**). Removal of CAN_THC_ elicited more active responses than VEH on extinction days 1-3 (interaction: *F*_(12,192)_=1.95, *p*=.03, 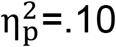; post-hoc *p’*s≤.03). Removal of cannabis increased inactive responding in all groups (*F*_(1,32)_=12.23, *p*=.001, 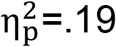). Removal of CAN_CBD_ elicited fewer inactive responses than removal of VEH on day 6 and fewer than removal of CAN_THC_ on day 4 (F_(12,192)_=3.12, *p*<.001, 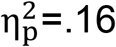, post-hoc *p*’s≤.04, **Fig. 4A**). Interestingly, removal of CAN_CBD_ increased the number of sessions rats needed to reach the extinction criterion compared to CAN_THC_ (*F*_(2,32)_=4.09, *p*=.03, 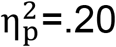, post-hoc *p*=.03, **Fig. 4B**).

**Figure 4.**
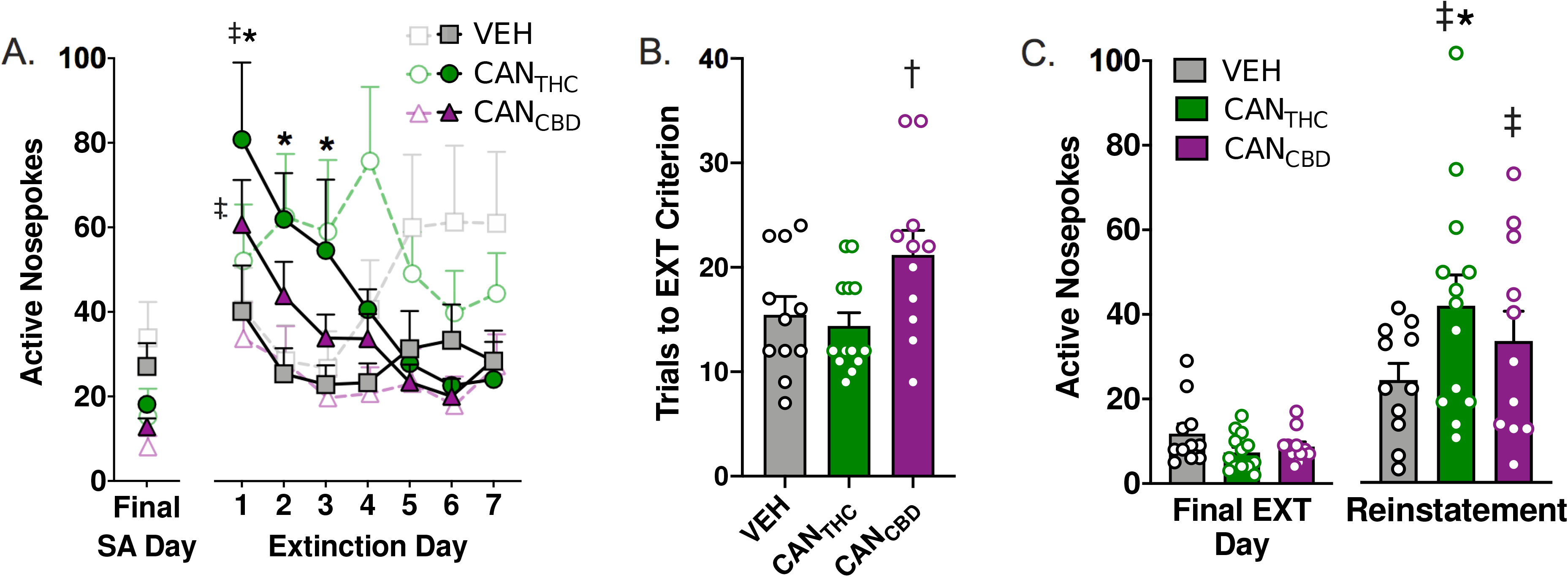
Cannabis vapor supports conditioned drug seeking in the absence of drug availability or in the presence of drug-related cues. **(A)** Active (colored symbols) and inactive (open symbols) responding for CAN_THC_, CAN_CBD_, or VEH vapor on the final day of self-administration training (left) and during the first 7 days of extinction training (right). **(B)** Number of trials required to meet extinction criterion (i.e., ≥ 50% decrease in active nosepoke responses relative to the final self-administration day during the final two extinction sessions). **(C)** Number of nosepoke responses made on the active operanda for CAN_THC_, CAN_CBD_, or VEH vapor on the final day of extinction (left) and during a cue-induced reinstatement test (left). n=11-13/group, *p* ≤ .05. ‡ denotes significant difference in responding relative to the final day of **(A)** self-administration or **(C)** extinction training. * denotes significant differences between CAN_THC_ and VEH groups. # denotes significant differences between CAN_THC_ and CAN_CBD_ groups. † denotes significant differences between CAN_CBD_ and VEH groups.

### CAN_THC_- or CAN_CBD_-Paired Stimuli Elicit Reinstatement of Cannabis Vapor-Seeking Behavior

Response-contingent presentation of either the CAN_THC_- or the CAN_CBD_-paired (but not the VEH-paired) light cue increased active responses at test relative to the last extinction training day (interaction: *F*(2,32)=3.48, *p*=.04, 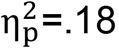; post-hoc *p*’s<.002, Fig. 4C). The CANTHC-paired cue elicited more active responses than the VEH-paired cue (*p*=.05) (Fig. 4C). Response-contingent cue presentation also altered inactive responding over the final extinction and reinstatement test days (*F*(2,32)=3.38, *p*=.047, 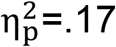), with CAN_CBD_ rats making fewer inactive responses than VEH rats irrespective of cue presentation (*p*=.04).

## DISCUSSION

Obstacles in establishing a model of cannabis use that closely mimics the human experience have limited our ability to study the neural mechanisms and impact of cannabis use [1,32]. In the current study, we present evidence supporting the feasibility of a novel preclinical model of cannabis self-administration that employs response-contingent delivery of vaporized cannabis extracts.

Our findings indicate that THC-dominant cannabis vapor (CAN_THC_) has reinforcing properties (**Fig. 1**). Despite variability in the rates of self-administration, both cannabis self-administering groups exhibited stable rates of responding under each schedule of reinforcement (**Fig. 1C**) and demonstrated robust discrimination between active and inactive operanda (**Fig. 1K**). The pattern of responding was consistent with a preserved loading dose phenomenon, with the majority of responding occurring during the first 15 min of the session (**Fig. 1D-F**). Importantly, active nosepoke responding was highest in the CAN_THC_ self-administering group, and only the CAN_THC_ self-administering group exhibited an increase in responding and maintained the number of vapor deliveries earned under FR-2 and FR-4 reinforcement schedules relative to the FR-1 schedule (**Fig. 1G**). Similarly, only the CAN_THC_ self-administering group exhibited higher break points and shorter latency to re-initiate responding post-vapor delivery under the PR reinforcement schedule relative to VEH rats (**Fig. 1M-N**).

Cannabis vapor also supported the acquisition of conditioned motivational effects by an initially neutral environmental stimulus (**Fig. 4**). Upon the removal of cannabis vapor reinforcement, both cannabis-trained groups increased responding relative to their final day of self-administration (i.e., extinction burst; **Fig. 4A**), but only the CAN_THC_-trained group made more responses than the VEH-trained control group. Similarly, only the CAN_THC_-trained group exhibited greater cue-induced reinstatement of cannabis-seeking behavior relative to the VEH-trained control group (**Fig. 4C**). These data indicate that vaporized cannabis extracts elicit goal-directed responding and cue-induced drug-seeking behavior, and the THC concentrations present in the extracts determine their reinforcing properties.

The validity of this model is further strengthened by our findings that response-contingent delivery of vaporized cannabis extracts produced behavioral and physiological changes that are consistent with those in human cannabis users (**Fig. 2**). The number of CAN_THC_ and CAN_CBD_ vapor deliveries obtained positively correlated with THC and CBD plasma concentrations immediately after the session (**Fig. 2A-B**). Self-administered CAN_THC_ reliably reduced locomotor activity during the session (**Fig. 2E**) and increased the duration of inactivity in the metabolic chambers during the first 2 hours following the self-administration session (**Fig. 2F**). Self-administered CAN_THC_ also increased daily food intake and energy expenditure (**Fig. 2G-H**). These data mirror observations of increased caloric intake and energy expenditure of chronic cannabis users [33–36]. Moreover, CB1R binding was significantly reduced in the hippocampus 24 hours after the final self-administration in CAN_THC_ rats compared to VEH rats (**Fig. 3A**), which is in line with studies in human cannabis users [19] and rodents repeatedly injected with THC [20–23]. Notably, biologically relevant THC and CBD concentrations were observed in brain tissue 24 hours after the final self-administration session (**Fig. 2C**), which may account for the lack of abstinence-induced anxiety-like behavior observed in this study (**Fig. S6**). Brain CBD concentrations were higher than brain THC concentrations (**Fig. 2D**), probably due to the long half-life of CBD (i.e., 24-48 hr) [37]. THC tissue concentration was low but detectable, and similar in both CAN_THC_ and CAN_CBD_ rats. CBD inhibits the cytochrome P450 family of liver enzymes that are primarily responsible for THC metabolism [38]. Thus, comparable brain THC concentrations following the CAN_THC_ and CAN_CBD_ regimens might reflect greater CBD-mediated inhibition of THC metabolism in the CAN_CBD_ group. This possibility will need to be systematically evaluated in future studies.

The vapor self-administration procedure likely facilitated volitional cannabis exposure. It has been well documented for other drugs of abuse that control over drug administration profoundly alters the subjective effects of drug intake, as well as the associated neurochemical responses [39–41]. Following passive intravenous THC administration, humans report aversive effects that are most often associated with the dose and infusion rate employed [17]. Thus, it is perhaps unsurprising that intravenous THC delivery generally fails to support self-administration in rodents (see [32] for review). The cannabis extracts used in the current study contain THC, CBD, and other naturally-occurring phytocannabinoids, and these phytocannabinoids may mitigate the aversive effects of THC [42]. Notably, CBD facilitates intravenous THC self-administration in rodents [12] (but see [43]) and offsets some of the pharmacological and behavioral effects of THC in humans [44] and rodents [9]. Furthermore, a passive THC+CBD vapor pre-exposure regimen facilitates the acquisition of intravenous THC+CBD self-administration, possibly by offsetting the novelty of THC intoxication or aversive stimulus properties prior to operant training [12]. Thus, the subjective and/or motivational effects of cannabis vapor delivery are different from effects of intravenous THC administration, and future studies should use whole cannabis preparations when possible and consider the route of administration when attempting to generalize effects to human cannabis users.

THC and CBD concentrations vary widely across commercially available cannabis products. As such, it will be important to study the extent to which modifying cannabinoid constituent concentrations alters the propensity for self-administration. In the current study, we compared responding for two different cannabis extracts: one extract containing 28.4% THC and 1.1% CBD (CAN_THC_) and another containing 1.96% THC and 59.3% CBD (CAN_CBD_). We observed key differences between the effects of CAN_THC_ and CAN_CBD_ extracts. First, CAN_THC_, but not CAN_CBD_, elicited significant reinforcing effects as indicated by augmented operant responding upon increases in schedule demand (**Fig. 1C**). Second, CAN_THC_ had greater motivational effects as indicated by higher breakpoints and a reduced latency to initiate responding following vapor delivery compared to CAN_CBD_ (**Fig. 1M-N**). Although both CAN_THC_ and CAN_CBD_ self-administration regimens were sufficient to increase drug seeking upon the removal of the reinforcer (i.e., extinction) or in the presence of vapor-associated cues (i.e., reinstatement) relative to responding during the last self-administration session and relative to the absence of cues, respectively, only the CAN_THC_ regimen elicited response rates above that of the VEH group (**Fig. 4A** and **4C**). Thus, CAN_THC_ vapor has greater reinforcing properties and stronger conditioned motivational effects than CAN_CBD_ and VEH vapor. Interestingly, the CAN_CBD_ regimen produced the strongest discrimination between active and inactive operanda despite reduced rates of responding. Furthermore, CAN_CBD_ elicited drug-seeking behavior that was more resistant to extinction, as indicated by a larger number of sessions required to reach extinction criterion relative to CAN_THC_ (**Fig. 4B**). Detectable brain concentrations of THC (**Fig. 2C**) and a CBD-mediated decrease in the aversive properties of THC likely reconcile these findings with studies indicating a lack of CBD-mediated reinforcement [45,46].

Altogether, findings from the present study strongly support the utility of a response-contingent vapor delivery protocol as a means to model stable and pharmacologically relevant levels of cannabis intoxication in human populations and further examine the neurobiological mechanisms of cannabis intake and cannabis-seeking behaviors. Ultimately, this model will permit finer interrogation of the effects of cannabis on the brain and behavior and help to identify causal factors that increase the susceptibility for developing cannabis use disorders.

## Supporting information

Cannabis Vapor Self-Administration - Progressive Ratio Responding

## REFERENCES

1. McLaughlin, RJ (2018): Toward a translationally relevant preclinical model of cannabis use. Neuropsychopharmacology. 43(1):213.

2. Laprairie RB, Bagher AM, Kelly ME, Dupré DJ, Denovan-Wright EM. (2014): Type 1 cannabinoid receptor ligands display functional selectivity in a cell culture model of striatal medium spiny projection neurons. J Biol Chem. 289(36):24845–62.

3. Fisar Z. (2009): Phytocannabinoids and endocannabinoids. Curr Drug Abuse Rev. 2(1):51–75.

4. Russo EB. (2011): Taming THC: potential cannabis synergy and phytocannabinoid-terpenoid entourage effects. Br J Pharmacol. 163(7):1344–64.

5. Hill AJ, Williams CM, Whalley BJ, Stephens GJ. (2012): Phytocannabinoids as novel therapeutic agents in CNS disorders. Pharmacol Ther. 133(1):79–97.

6. Lewis MA, Russ EB, Smith KM. (2018): Pharmacological foundations of cannabis chemovars. Planta Med. 84(4):225–233.

7. Laprairie RB, Bagher AM, Kelly ME, Denovan-Wright EM. (2015): Cannabidiol is a negative allosteric modulator of the cannabinoid CB1 receptor. Br J Pharmacol. 172(20):4790–805.

8. Laprairie RB, Bagher AM, Kelly ME, Denovan-Wright EM. (2016): Biased Type 1 Cannabinoid Receptor Signaling Influences Neuronal Viability in a Cell Culture Model of Huntington Disease. Mol Pharmacol. 89(3):364–75.

9. Englund A, Morrison PD, Nottage J, Hague D, Kane F, Bonaccorso S, et al. (2013): Cannabidiol inhibits THC-elicited paranoid symptoms and hippocampal-dependent memory impairment. J Psychopharmacol. 27(1):19–27.

10. Greene NZ, Wiley JL, Yu Z, Clowers BH, Craft RM. (2018): Cannabidiol modulation of antinociceptive tolerance to ∆_9_-tetrahydrocannabinol. Psychopharmacology (Berl). 235(11):3289–3302.

11. Tanda G. (2016): Preclinical studies on the reinforcing effects of cannabinoids. A tribute to the scientific research of Dr. Steve Goldberg. Psychopharmacology (Berl). 233(10):1845–1866.

12. Spencer S, Neuhofer D, Chioma VC, Garcia-Keller C, Schwartz DJ, Allen N, et al. (2018): A Model of Δ9-Tetrahydrocannabinol Self-administration and Reinstatement That Alters Synaptic Plasticity in Nucleus Accumbens. Biol Psychiatry. 84(8): 601–610.

13. Sexton M, Cuttler C, Finnell JS, Mischley LK. (2016): A Cross-Sectional Survey of Medical Cannabis Users: Patterns of Use and Perceived Efficacy. Cannabis Cannabinoid Res. 1(1):131–138.

14. Grotenhermen F. (2003): Pharmacokinetics and pharmacodynamics of cannabinoids. Clin Pharmacokinet. 42(4):327–60.

15. Huestis MA. (2007): Human cannabinoid pharmacokinetics. Chem Biodivers. 4(8):1770–804.

16. Hložek T, Uttl L, Kadeřábek L, Balíková M, Lhotková E, Horsley RR, et al. (2017): Pharmacokinetic and behavioural profile of THC, CBD, and THC+CBD combination after pulmonary, oral, and subcutaneous administration in rats and confirmation of conversion in vivo of CBD to THC. Eur Neuropsychopharmacol. 27(12):1223–1237.

17. Carbuto M, Sewell RA, Williams A, Forselius-Bielen K, Braley G, Elander J. (2012): The safety of studies with intravenous Δ^9^-tetrahydrocannabinol in humans, with case histories. Psychopharmacology (Berl). 219(3):885–96.

18. Manwell LA, Charchoglyan A, Brewer D, Matthews BA, Heipel H, Mallet PE. (2014): A vapourized Δ(9)-tetrahydrocannabinol (Δ(9)-THC) delivery system part I: development and validation of a pulmonary cannabinoid route of exposure for experimental pharmacology studies in rodents. J Pharmacol Toxicol Methods. 70(1):120–7.

19. Ceccarini J, Kuepper R, Kemels D, van Os J, Henquet C, Van Laere K. (2015): [18F]MK-9470 PET measurement of cannabinoid CB1 receptor availability in chronic cannabis users. Addict Biol. 20(2): 357–67.

20. Romero J, Garcia-Palomero E, Castro JG, Garcia-Gil L, Ramos JA, Fernandez-Ruiz JJ. (1997): Effects of chronic exposure to delta-9-tetrahydrocannabinol on cannabinoid receptor binding and mRNA in several rat brain regions. Brain Res Mol Brain Res. 46(1-2): 100–8.

21. Silva L, Harte-Hargrove L, Izenwasser S, Frank A, Wade D, Dow-Edwards D. (2015): Sex-specific alterations in hippocampal cannabinoid 1 receptor expression following adolescent delta-9-tetrahydrocannabinol treatment in the rat. Neurosci Lett. 602: 89–94.

22. Farquhar CE, Breivogel CS, Gamage TF, Gay EA, Thomas BF, Craft RM, Wiley JL. (2019): Sex, THC, and hormones: Effects on density and sensitivity of CB1 cannabinoid receptors in rats. Drug Alcohol Depend. 194: 20–27.

23. Kruse LC, Cao JK, Viray K, Stella N, Clark JJ. (2019): Voluntary oral consumption of Δ^9^-tetrahydrocannabinol by adolescent rats impairs reward-predictive cue behaviors in adulthood. Neuropsychopharmacology. 44(8): 1406–14.

24. Hasin DS, Keyes KM, Alderson D, Wang S, Aharonovich E, Grant BF. (2008): Cannabis withdrawal in the United States: results from NESARC. J Clin Psychiatry. 69(9):1354–63.

25. Nguyen JD, Aarde SM, Vandewater SA, Grant Y, Stouffer DG, Parsons LH, et al. (2016): Inhaled delivery of Δ(9)-tetrahydrocannabinol (THC) to rats by e-cigarette vapor technology. Neuropharmacology. 109:112–120.

26. Javadi-Paydar M, Nguyen JD, Kerr TM, Grant Y, Vandewater SA, Cole M, Taffe MA. (2018): Effects of Δ9-THC and cannabidiol vapor inhalation in male and female rats. Psychopharmacology (Berl). 235(9):2541–2557.

27. Walker BM, Koob GF. (2007): The gamma-aminobutyric acid-B receptor agonist baclofen attenuates responding for ethanol in ethanol-dependent rats. Alcohol Clin Exp Res. 31(1):11–18.

28. Britch SC, Wiley JL, Yu Z, Clowers BH, Craft RM. (2017): Cannabidiol-Δ9-tetrahydrocannabinol interactions on acute pain and locomotor activity. Drug Alcohol Depend. 175:187–197.

29. Lee TT, Hill MN. (2013): Age of stress exposure modulates the immediate and sustained effects of repeated stress on corticolimbic cannabinoid CB1 receptor binding in male rats. Neuroscience. 249: 106–14.

30. Berger AL, Henricks AM, Lugo JM, Wright HR, Warrick CR, Sticht MA, Morena M, et al. (2018): The lateral habenula directs coping styles under conditions of stress via recruitment of the endocannabinoid system. Biol Psychiatry. 84(8): 611–23.

31. Henricks AM, Berger AL, Lugo JM, Baxter-Potter LN, Bieniasz KV, Petrie G, et al. (2017): Sex- and hormone-dependent alterations in alcohol withdrawal-induced anxiety and corticolimbic endocannabinoid signaling. Neuropharmacology. 124:121–133.

32. Melis M, Frau R, Kalivas PW, Spencer S, Chloma V, Zamberletti E, et al. (2017): New vistas on cannabis use disorder. Neuropharmacology. 124:62–72.

33. Smit E, Crespo CJ. (2001): Dietary intake and nutritional status of US adult marijuana users: results from the Third National Health and Nutrition Examination Survey. Public Health Nutr. 4(3):781–786.

34. Rodondi N, Pletcher MJ, Liu K, Hulley SB, Sidney S, Coronary Artery Risk Development in Young Adults (CARDI) Study. (2006): Marijuana use, diet, body mass index, and cardiovascular risk factors (from the CARDIA study). Am J Cardiol. 98(4):478–484.

35. Le Strat Y, Le Foll B. (2011): Obesity and cannabis use: results from 2 representative national surveys. Am J Epidemiol. 174(8):929–933.

36. Meier MH, Pardini D, Beardslee J, Matthews KA. (2019): Associations between cannabis use and cardiometabolic risk factors: A longitudinal study of men. Psychosom Med. 81(3):281–288.

37. Millar SA, Stone NL, Yates AS, O’Sullivan SE. (2018): A systematic review on the pharmacokinetics of cannabidiol in humans. Front Pharmacol. 9:1365.

38. Zendulka O, Dovrtělová G, Nosková K, Turjap M, Šulcová A, Hanuš L, Juřica J. (2016): Cannabinoids and cytochrome P450 interactions. Curr Drug Metab. 17(3):206–226.

39. Stefanski R, Ladenheim B, Lee SH, Cadet JL, Goldberg SR. (1999): Neuroadaptations in the dopaminergic system after active self-administration but not after passive administration of methamphetamine. Eur J Pharmacol. 371(2-3):123–35.

40. Donny EC, Caggiula AR, Rose C, Jacobs KS, Mielke MM, Sved AF. (2000): Differential effects of response-contingent and response-independent nicotine in rats. Eur J Pharmacol. 402(3):231–40.

41. Stefański R, Ziółkowska B, Kuśmider M, Mierzejewski P, Wyszogrodzka E, Kołomańska P, et al. (2007): Active versus passive cocaine administration: differences in the neuroadaptive changes in the brain dopaminergic system. Brain Res. 1157:1–10.

42. Russo E, Guy GW. (2006): A tale of two cannabinoids: the therapeutic rationale for combining tetrahydrocannabinol and cannabidiol. Med Hypotheses. 66(2):234–246.

43. Wakeford AGP, Wetzell BB, Pomfrey RL, Clasen MM, Taylor WW, Hempel BJ, Riley AL. (2017): The effects of cannabidiol (CBD) on Δ-tetrahydrocannabinol (THC) self-administration in male and female Long-Evans rats. Exp Clin Psychopharmacol. 25(4):242–248.

44. Morgan CJ, Freeman TP, Schafer GL, Curran HV. (2010): Cannabidiol attenuates the appetitive effects of Delta 9-tetrahydrocannabinol in humans smoking their chosen cannabis. Neuropsychopharmacology. 35(9):1879–85.

45. Haney M, Malcolm RJ, Babalonis S, Nuzzo PA, Cooper ZD, Bedi G, et al. (2016): Oral cannabidiol does not alter the subjective, reinforcing or cardiovascular effects of smoked cannabis. Neuropsychopharmacology. 41(8):1974–82.

46. Viudez-Martinez A, Garcia-Gutierrez MS, Medrano-Relinque J, Navarron CM, Navarrete F, Manzanares J. (2019): Cannabidiol does not display drug abuse potential in mice behavior. Acta Pharmacol Sin. 40(3):358–364.

47. Fuchs RA, Higginbotham, JA, Hansen EJ. (2018): Animal models of drug addiction. In: Neural Mechanisms of Addiction (Ed.: Torregrossa, M.), Academic Press, Elsevier, pp. 3–22.

