## Supplemental Information for "Vaporized cannabis extracts have reinforcing properties and support conditioned drug-seeking behavior in rats"

**CONTENTS**

Supplemental Methods

**Figure S1.** Metabolic phenotype of rats self-administering $CAN_{THC}$, $CAN_{CBD}$, or VEH vapor.

**Figure S2.** Self-administration data for rats receiving systemic AM251 injections.

**Figure S3.** Effects of systemic AM251 treatment on instrumental responding for $CAN_{THC}$ or $CAN_{CBD}$ vapor (raw values).

**Figure S4.** Self-administration data for rats in the extinction/reinstatement experiment.

**Figure S5.** Inactive responding during extinction and cue-induced reinstatement.

**Figure S6.** Elevated plus maze behavior following forced abstinence from cannabis vapor.

Supplemental References

**SUPPLEMENTAL METHODS**

***Vapor Chamber Apparatus***

Eight 13.5” x 9.0” x 8.25” (L x W x H) 16.4 L vapor self-administration chambers (La Jolla Alcohol Research Inc.) were programmed using MED-Associates IV software to deliver response-contingent puffs of vapor. An uninterrupted unidirectional flow of air entered through a port in the front of the chamber, and air was removed by vacuum in the rear of the chamber lid. The air intake port pulled air through tubing connected to an air flow meter and tubing connected to a commercial e-cigarette cartridge (first generation: Protank 3 Dual Coil; 2.2 W coils; Kanger Tech, Shenzhen, China; second generation: SMOK Tank Baby Beast TFV8 with 0.2Ω M2 atomizer) filled with $CAN_{THC}$, $CAN_{CBD}$, or VEH. Experiments involving radiotelemetry, extinction, cue-induced reinstatement, CB1R antagonism, and quantification of plasma cannabinoid concentrations were conducted using first-generation vaporizers that delivered 10 s puffs of vapor. All other experiments were conducted using second-generation vaporizers that delivered 3 s puffs. These puff durations were chosen to match the volume of vapor produced by each system during response-contingent delivery. Two nosepoke operanda and associated cue lights were located on the rear wall of the chamber. The cartridge was connected to a vaporizer box (La Jolla Alcohol Research Inc.), and vapor puffs were delivered through the air intake port. Chamber air was evacuated through an activated charcoal filter (Carbatrol Corporation; Bridgeport, CT) or in-line Whatman HEPA-Cap filters (Millipore-Sigma, St. Louis, MI).

***Plasma Cannabinoid Concentrations***

An ultra-performance liquid chromatography system (Waters Acquity I-Class UPLC, Milford, MA, USA) coupled with a quadrupole time of flight mass spectrometer (QTOF, Waters Xevo G2, Manchester, UK) was used to quantify the concentration of THC and CBD in plasma using the method described in Greene et al., 2018. Briefly, samples were centrifuged at 8000 rpm for 10 min and 185 uL of the resulting supernatant was spiked with 15 uL of solution containing 200 ppb each of the THC-d3 and CBD-d3 deuterated standards (Cerilliant, Round Rock, TX). After the internal standard was added, protein precipitation was promoted by adding 400 uL of cold acetonitrile dropwise while vortexing. Samples were centrifuged at 4000 g for 10 min at 25°C. 0.6 mL of 1% ammonium hydroxide was added to the sample and vortexed before solid phase extraction (SPE). A mixed mode SPE cartridge (OAXIS Max 1 cc, Waters, Ireland) was used for cannabinoid isolation. Each SPE cartridge was conditioned with 1 mL of methanol followed by 1 mL of 1% ammonium hydroxide. Conditioned samples were then loaded onto the SPE cartridge and pulled through the system using a light vacuum (~1–2 psi). 35% acetonitrile (0.5 mL) was added and allowed to dry under full vacuum for 10 min. Samples were eluted with 1.5 mL of a hexane/ethyl acetate/acetic acid (49:49:2, v/v/v). The eluent was then evaporated under nitrogen at room temperature and 100 uL of a methanol:water solution (80:20, v/v) was transferred to an autosampler vial. 50-mm C18 BEH UPLC columns (Waters, Milford, MA, USA) were kept at 40°C. Mobile phases were (A) high purity water (Fisher Scientific Co., Fair Lawn, NJ) with 0.1% formic acid and (B) pure acetonitrile (Fisher Scientific Co., Fair Lawn, NJ) with 0.1% formic acid. A total of 10 uL of each prepared sample was injected onto the column. Data were then analyzed using TargetLynx (Waters, Milford, MA) with the following parameters: Retention time window: ± 0.2 min, Response use: integrated area, Polynomial Type: linear, Weighting function: 1/X.

***Tissue Cannabinoid Concentrations***

Extraction and quantification of THC, CBD, THC-COOH and 11-OH-THC were carried out as previously described (Baglot et al., under review). Briefly, frozen brain tissue was briefly weighed and then manually homogenized (with a glass rod) in borosilicate glass culture tubes containing 2ml of acetonitrile with 1ng of each THC-d3, CBD-d3, THC-COOHd3 and 11-OH-THCd3. Samples were then sonicated for 30min in an ice bath and incubated overnight at -20C to precipitate proteins. The following day samples were centrifuged at 1500xg to remove particulates. The supernatant from each sample was transferred to a new glass tube and evaporated under nitrogen, the tube was then washed once with 350ul acetonitrile (to recapture any lipids adhering to the glass wall) and the acetonitrile was dried under nitrogen gas again. After completely drying, the samples were re-suspended in 200ul of 1:1 methanol:water and stored at -80C until analysis by liquid chromatography mass spectrometry. Quantification of these molecules using mass spectrometry was performed using an Eksigent Micro LC200 coupled with an AB Sciex QTRAP 5500 mass spectrometry (AB Sciex, Ontario, Canada) as previously described (Baglot et al., under review). The data were acquired in positive electrospray ionization (ESI) and multiple reaction monitoring (MRM) mode and amount of each molecule was normalized to frozen tissue weight.

***Radiotelemetry Recordings***

Rats were anesthetized with an isoflurane/oxygen vapor mixture (isoflurane 5% induction, 1-3% maintenance) and a 1-cm midline vertical incision was made inferior to the xyphoid space. Sterile radiotelemetry transmitters (Starr Life Sciences Corp., Oakmont, PA; PTD 4000) were inserted under the muscular layer and sutured in place using absorbable 4-0 silk sutures. The muscle layer was closed with 5-0 vicryl suture, and the skin was closed with non-absorbable wax-coated 4-0 silk suture. Rats received meloxicam (2 mg/kg, sc) for 3 days for post-operative pain management. After at least five days of recovery, rats were randomly assigned to treatment groups and trained to nosepoke for $CAN_{THC}$, $CAN_{CBD}$, or VEH vapor. Radiotelemetry transmissions indexing locomotor activity and body temperature were collected daily during one-hour self-administration sessions by Respironics ER-4000 receiver plates placed under the chambers.

***Metabolic Phenotyping***

A subset of rats were housed individually in metabolic cages (dimensions = L 18“ x W 9.5” x H 8.1”) (Promethion, Sable Systems International) throughout self-administration training, during which feeding behavior, water intake, energy expenditure, physical activity and respiratory quotient (RQ) were monitored. Cages included a ceiling-mounted food hopper (3-mg resolution) and a water spigot connected to load cells (MM-1, Sable Systems International) for food and water intake monitoring, respectively. Ad *libitum* access to the food hopper and water were allowed throughout the study. X- and Y-axis (horizontal plane) photoelectric beam motion detectors were positioned around each cage to assess ambulatory activity. In addition to total distance travelled, the total time spent engaging in lounging behavior was quantified and sub-divided into short and long lounging. Short lounging was defined as any period of inactivity greater than 15 s but less than 60 s, whereas long lounging was defined as any period of inactivity greater than 60 s.

Respiratory gases were measured with an integrated fuel cell oxygen analyzer, spectrophotometric

$CO_2$ analyzer, and capacitive water vapor partial pressure analyzer (GA3m1, Sable Systems International). The Promethion system uses a pull-mode, negative pressure system. The multi-channel mass flow generator measures and controls air flow (FR8-1, Sable Systems International). The incurrent flow rate was set at 2000 mL/min. Water vapor was continuously measured and its dilution effect on $O_2$ and $CO_2$ was compensated for mathematically in the analysis (Lighton and Turner, 2008). Respiratory exchange quotient (RQ) was calculated as the ratio of $CO_2$ production to $O_2$ consumption. Energy expenditure was calculated using the Weir equation: $Kcal/h = 60 * (0.003941 * VO_2 + 0.001106 * VCO_2)$ (Weir, 1949). Data were acquired using MetaScreen v. 2.2.8, the raw data obtained were processed using ExpeData v. 1.8.2 (Sable Systems International), and Macros 10 and 13 were used for data organization and transformation.

Values were collected and averaged over the final 10 days of self-administration, as this corresponds to the maintenance phase of drug seeking. Each day, rats were weighed, transferred to individual holding cages, and transported to the vapor self-administration system between 11h00 and 14h00. Daily values are averaged for active and inactive phases excluding the period encompassing transportation and self-administration training.

***Elevated Plus Maze Test***

The EPM apparatus consisted of a raised Plexiglas platform (28.5 inches high) with two open exposed arms and two darker enclosed arms of equal length (21.5 inches/arm) (Med Associates Inc., St. Albans, VT). The floors were made of clear Plexiglas and the walls of both closed arms were black. Twenty-four hours after their final self-administration session, rats were individually placed in the center of the maze and allowed to freely explore the maze for 5 min. All tests were run in dim lighting (~10 lux), and behaviors were recorded with Noldus Ethovision XT behavioral tracking software. The number of entries and percent time spent in the open and closed arms, and distance traveled in the open arms of the EPM were compared across groups. The frequency of risk assessment behaviors (i.e., head-dips, stretch-attend postures, and rearing) was also scored manually by trained research assistants blinded to treatment conditions.

**SUPPLEMENTAL RESULTS**

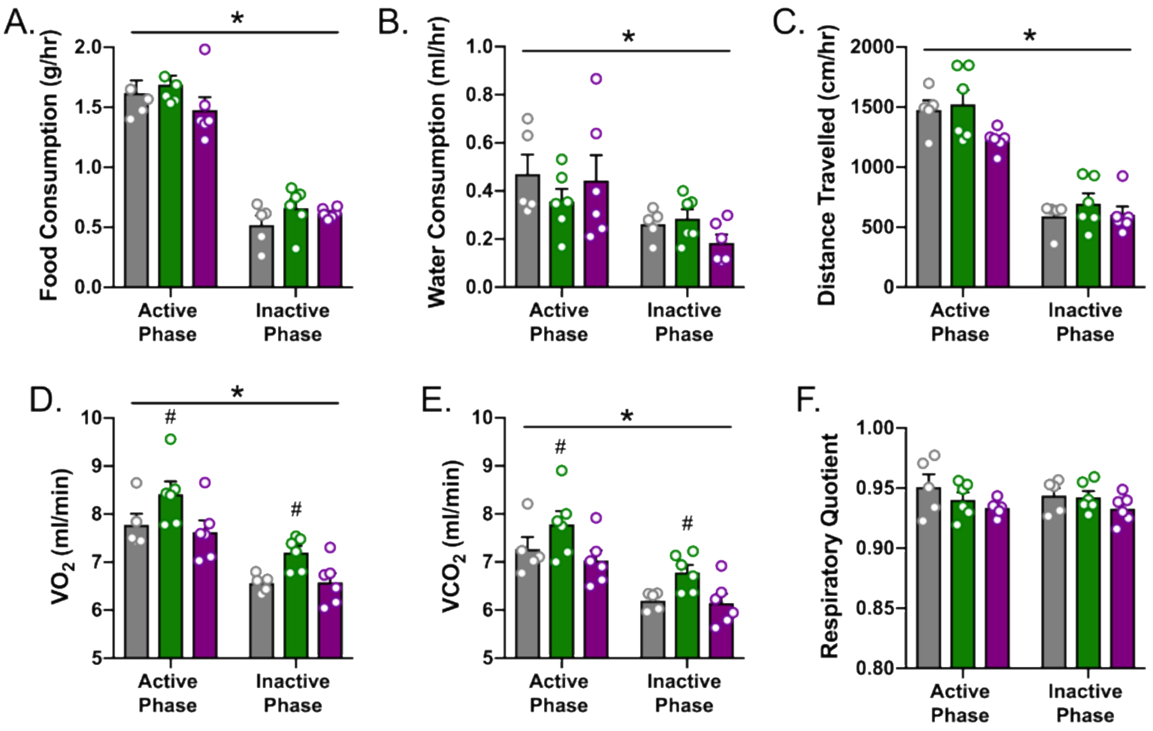


**Figure S1. Metabolic phenotype of rats self-administering CAN$_{THC}$, CAN$_{CBD}$, or VEH vapor.** Self-administration of CAN$_{THC}$ or CAN$_{CBD}$ did not significantly alter the rate of **(A)** food consumption, **(B)** water consumption, or **(C)** distance travelled compared to VEH when tabulated over the active and inactive phases. The rate of **(D)** $O_2$ consumption ($VO_2$) and **(E)** $CO_2$ production ($VCO_2$) was significantly increased among rats self-administering CAN$_{THC}$ vapor relative to rats in the CAN$_{CBD}$ condition. As expected, rates of food and water consumption, distance travelled, $VO_2$, and $VCO_2$ were greater during the active phase relative to the inactive phase **(A-E)**. **(F)** The respiratory quotient did not significantly differ by phase or treatment group. Significant effects of phase are denoted by * and significant differences between CAN$_{THC}$ and CAN$_{CBD}$ groups are denoted by #. n=5-6/group, $p \leq .05$.

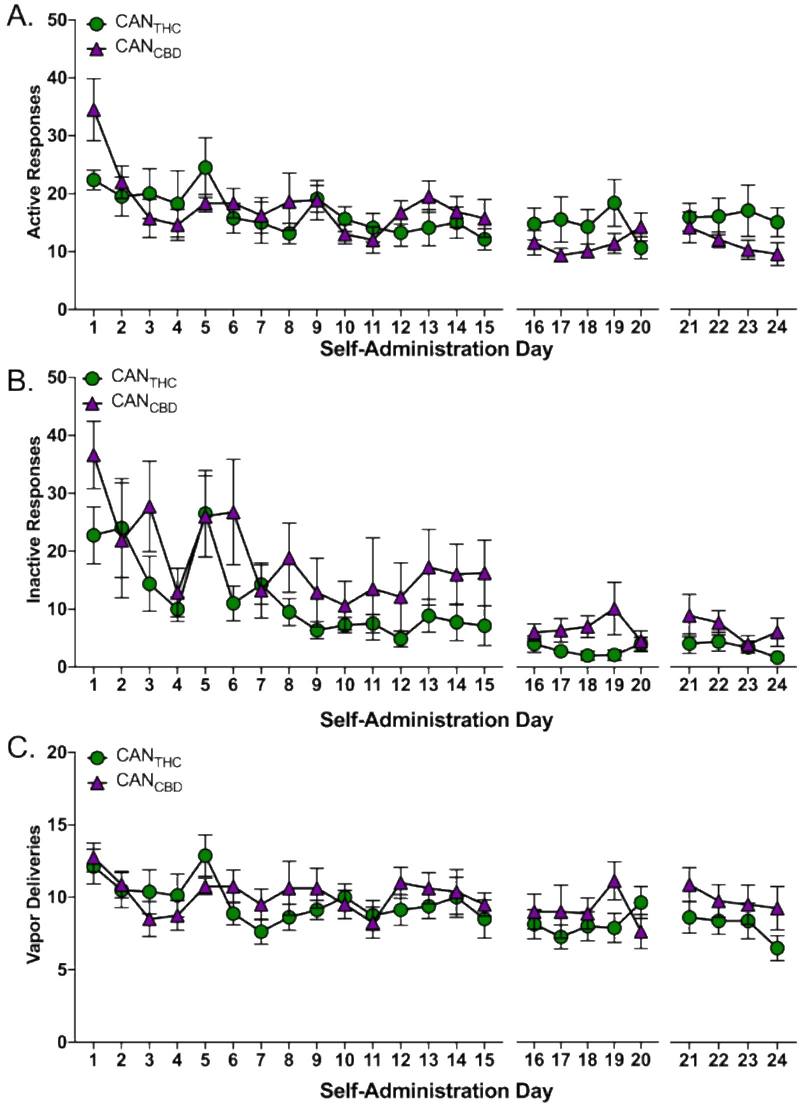


**Figure S2. Self-administration data for rats receiving systemic AM251 injections.** The effect of systemic CB1R antagonism with AM251 (0, 1, or 3 mg/kg, ip) on rates of **(A)** active nosepoke responding, **(B)** inactive nosepoke responding, and **(C)** the number of vapor deliveries earned during daily 1 hr sessions. There were no significant differences between active responding, inactive responding, or vapor deliveries earned in rats trained to self-administer $CAN_{THC}$ or $CAN_{CBD}$ vapor. Data are presented across self-administration days and gaps between days 15-16 and 20-21 indicate days on which AM251 injections occurred. Mock injections were administered on the two days preceding an AM251 challenge. n=8/group, $p \leq .05$.

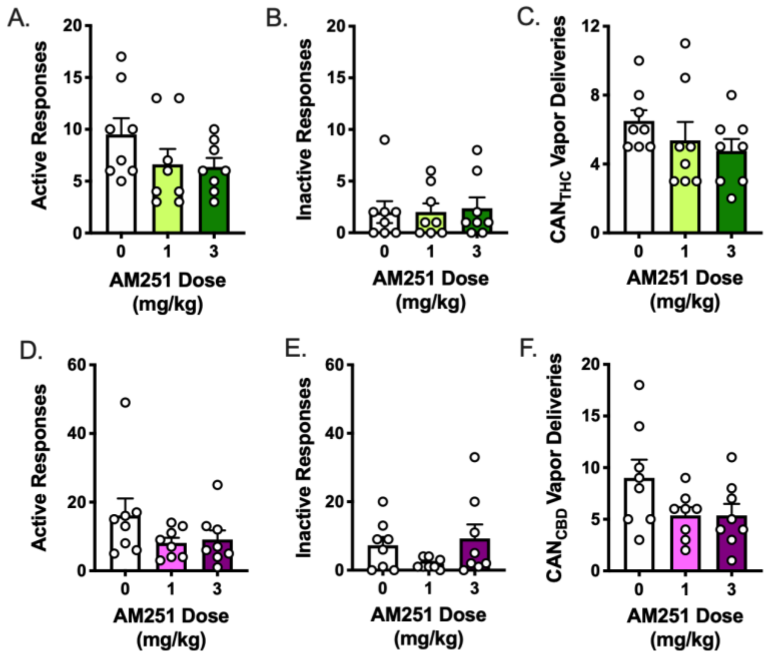


**Figure S3. Effects of systemic AM251 treatment on instrumental responding for $CAN_{THC}$ or $CAN_{CBD}$ vapor (raw values). (A)** There was a trend for administration of AM251 to decrease the total number of active nosepoke responses made for $CAN_{THC}$ vapor ($F_{(2,14)}$ = 2.87, *p* = .09, $\eta_{p2}$=.31), but post-hoc analyses did not indicate significant group differences (0 mg/kg vs. 1 mg/kg: *p* = .19; 0 mg/kg vs. 3 mg/kg: *p* = .07). There was no significant effect of AM251 administration on **(B)** the total number of inactive nosepoke responses made for $CAN_{THC}$ vapor ($F_{(2,14)}$ = 0.05, *p* = .93) or **(C)** the number of $CAN_{THC}$ vapor deliveries earned ($F_{(2,14)}$ = 2.75, *p* = .12, $\eta_{p2}$=.31). There was no effect of AM251 treatment on the total number of **(D)** active nosepoke responses for $CAN_{CBD}$ vapor ($F_{(2,14)}$ = 1.60, *p* = .25), **(E)** inactive nosepoke responses for $CAN_{CBD}$ vapor ($F_{(2,14)}$ = 1.98, *p* = .19), or **(F)** $CAN_{CBD}$ vapor deliveries earned ($F_{(2,14)}$ = 2.38, *p* = .16). *n* = 8/group.

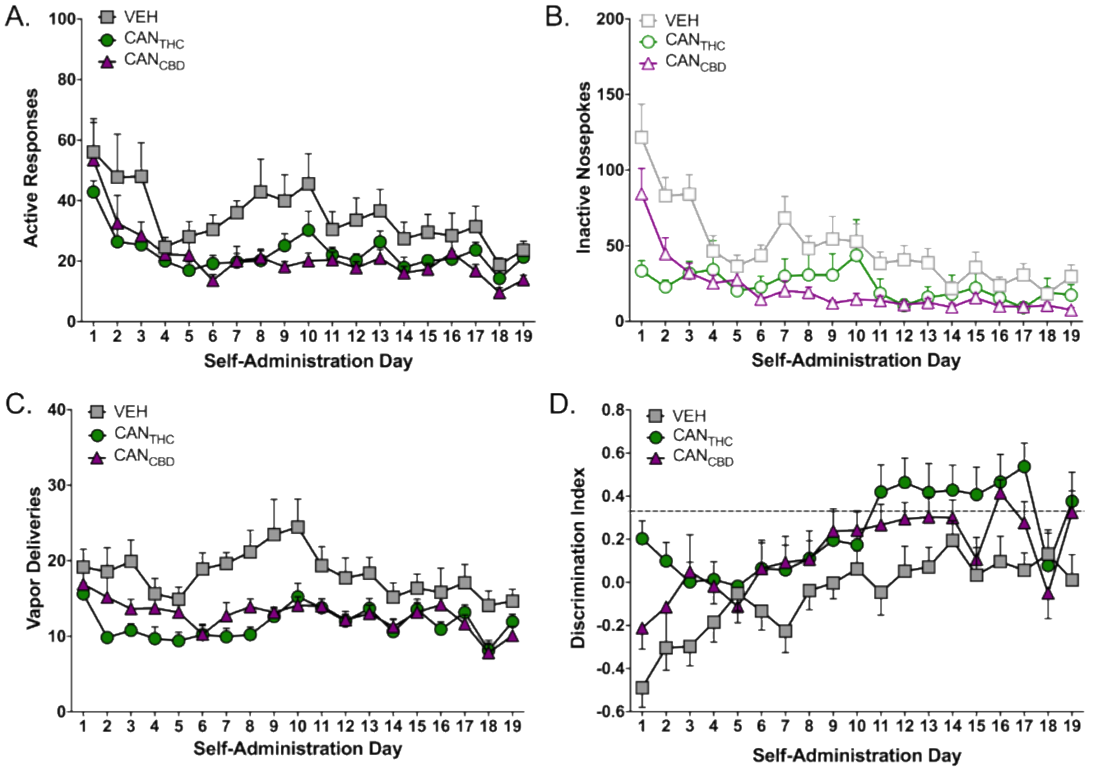


**Figure S4. Self-administration data for rats in the extinction/reinstatement experiment.** During vapor self-administration training, active nosepoke responding **(A)** declined and stabilized over time (time main effect, $F_{(18,576)}$ = 9.82, $p$ = .0001, $\eta_{p2}$= .23), and CAN$_{THC}$ or CAN$_{CBD}$ reinforcement elicited less responding relative to VEH overall (group main effect, $F_{(2,32)}$ = 4.55, $p$ = .01, $\eta_{p2}$=.22; $p$'s ≤ .04; interaction, *ns*). Group differences in inactive nosepoke responding **(B)** varied across time (interaction, $F_{(36,576)}$ = 6.50, $p$ < .001, $\eta_{p2}$=.28; main effects not shown). Specifically, CAN$_{THC}$ reinforcement elicited fewer inactive nosepoke responses relative to VEH on days 1-3 and 7 ($p$'s ≤ .001) and relative to CAN$_{CBD}$ on day 1 ($p$ = .001). Furthermore, CAN$_{CBD}$ reinforcement elicited fewer inactive nosepoke responses than VEH on days 1-3, 7, and 9-10 ($p$'s ≤ .02). The number of vapor deliveries obtained **(C)** fluctuated before stabilizing over time (time main effect, $F_{(18,576)}$ = 4.79, $p$ = .0001, $\eta_{p2}$= .13), and the groups obtained fewer CAN$_{THC}$ or CAN$_{CBD}$ vapor deliveries than VEH vapor deliveries (group main effect, $F_{(2,32)}$ = 8.57, $p$ = .001, $\eta_{p2}$ = .34; $p$'s ≤ .01; interaction, *ns*). (**D**) Group differences in discrimination indices also varied across time (interaction, $F_{(36,576)}$ = 1.50, $p$ = .03, $\eta_{p2}$ = .08; main effects not shown). CAN$_{THC}$ produced greater discrimination between active and inactive operanda relative to VEH on days 1-2, 11-12, 15, and 17 ($p$'s ≤ .04) and relative to CAN$_{CBD}$ on day 1 ($p$ = .02). $n$ = 11–13/group.

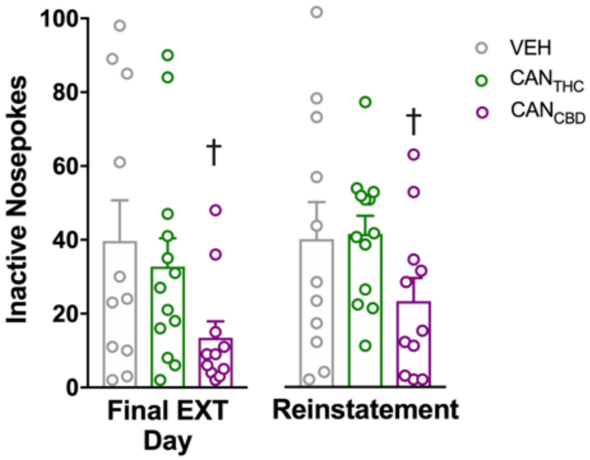


**Figure S5. Inactive responding during extinction and cue-induced reinstatement.** There was a main effect of treatment on inactive responding when tabulated over the final extinction day and reinstatement test day ($F_{(2,32)}$=3.38, *p*=.047, $\eta_{p2}$=.17). Specifically, CAN$_{CBD}$ rats made fewer inactive responses than VEH rats, irrespective of cue presentation (*p*=.043). † denotes significant differences between CAN$_{CBD}$ and VEH groups at *p*<.05.

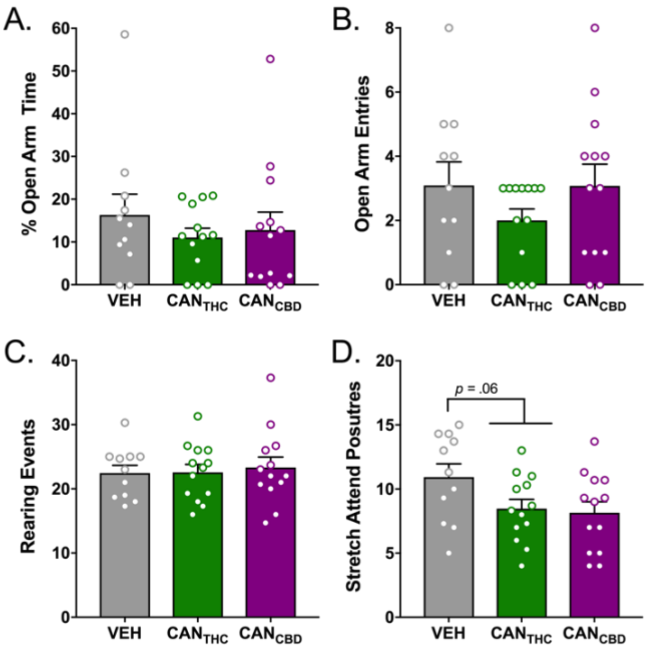


**Figure S6. Elevated plus maze behavior following forced abstinence from cannabis vapor.** When EPM behavior was measured 24 hr after the final vapor self-administration session, neither $CAN_{THC}$ nor $CAN_{CBD}$ history altered **(A)** the percentage of time spent in the open arms of the EPM ($F_{(2,34)}$=0.53, $p$=.48, $\eta_{p2}$=.03), **(B)** the number of open arm entries made ($F_{(2,34)}$=1.12, $p$=.34, $\eta_{p2}$=.06), **(C)** the number of rearing events ($F_{(2,34)}$=.12, $p$=.89, $\eta_{p2}$=.007), or **(D)** the frequency of stretch-attend postures ($F_{(2,34)}$=2.90, $p$=.07, $\eta_{p2}$=.15). However, there was a trend ($p$=.06) for $CAN_{THC}$ and $CAN_{CBD}$ rats to make fewer stretch-attend postures than VEH rats.

**SUPPLEMENTAL REFERENCES**

Baglot SL, Zhuo R, Parker LA, Rho J, McLaughlin RJ, Brechenmacher L, Hill MN (2019): Pharmacokinetic Comparison of Brain and Blood Levels of Delta-9-tetrahydrocannabinol and Metabolites Following Parenteral or Pulmonary Administration in Rats. Submitted.

Greene NZ, Wiley JL, Yu Z, Clowers BH, Craft RM. (2018): Cannabidiol modulation of antinociceptive tolerance to $\Delta_9$-tetrahydrocannabinol. *Psychopharmacology (Berl).* 235(11):3289-3302.

Lighton JR, Turner RJ. (2008): The hygric hypothesis does not hold water: abolition of discontinuous gas exchange cycles does not affect water loss in the ant Camponotus vicinus. *J Exp Biol.* 211(Pt 4):563-567.

Weir JB. (1949): New methods for calculating metabolic rate with special reference to protein metabolism. *J Physiol.* 109(1-2):1-9.
